# Visual mismatch responses index surprise signalling but not expectation suppression

**DOI:** 10.1101/2020.06.23.168187

**Authors:** Daniel Feuerriegel, Jane Yook, Genevieve L. Quek, Hinze Hogendoorn, Stefan Bode

## Abstract

The ability to distinguish between commonplace and unusual sensory events is critical for efficient learning and adaptive behaviour. This has been investigated using oddball designs in which sequences of often-appearing (i.e. expected) stimuli are interspersed with rare (i.e. surprising) deviants. Resulting differences in electrophysiological responses following surprising compared to expected stimuli are known as visual mismatch responses (VMRs). VMRs are thought to index co-occurring contributions of stimulus repetition effects, expectation suppression (that occurs when one’s expectations are fulfilled), and expectation violation (i.e. surprise) responses; however, these different effects have been conflated in existing oddball designs. To better isolate and quantify effects of expectation suppression and surprise, we adapted an oddball design based on Fast Periodic Visual Stimulation (FPVS) that controls for stimulus repetition effects. We recorded electroencephalography (EEG) while participants (N=48) viewed stimulation sequences in which a single face identity was periodically presented at 6 Hz. Critically, one of two different face identities (termed oddballs) appeared as every 7th image throughout the sequence. The presentation probabilities of each oddball image within a sequence varied between 10-90%, such that participants could form expectations about which oddball face identity was more likely to appear within each sequence. We also included ‘expectation neutral’ 50% probability sequences, whereby consistently biased expectations would not be formed for either oddball face identity. We found that VMRs indexed surprise responses, and effects of expectation suppression were absent. That is, ERPs were more negative-going at occipitoparietal electrodes for surprising compared to neutral oddballs, but did not differ between expected and neutral oddballs. Surprising oddball-evoked ERPs were also highly similar across the 10-40% appearance probability conditions. Our findings indicate that VMRs which are not accounted for by repetition effects are best described as an all-or-none surprise response, rather than a minimisation of prediction error responses associated with expectation suppression.

**Highlights:**

- We used a recently-developed oddball design that controls for repetition effects
- We found effects of surprise but not expectation suppression on ERPs
- Surprise responses did not vary by stimulus appearance probability

## 1. Introduction

We are highly adept at learning recurring patterns and statistical regularities that occur in our sensory environments, and we can exploit this knowledge to discriminate between commonplace and unusual sensory events. The computations that underpin this capability likely rely on statistics learned over time-scales ranging from hundreds of milliseconds to several minutes (Ulanovsky et al., 2004; Maheu et al., 2019). Characterising the computations that occur within different brain regions can help us understand how we generate and evaluate internal models of our environment, and how these models are used to facilitate detection of potential rewards (Schultz, 2016) and enact adaptive adjustments to decision-making strategies (Wessel, 2017; Wessel & Huber, 2019).

Detection of novel or unusual events has been investigated using visual oddball designs in which a stimulus is presented in high and low probability contexts. For example, in the high probability context, a critical stimulus A is presented frequently as the ‘standard’ stimulus, interspersed with a rare ‘deviant’ stimulus B (e.g., AABAAAAAAABAAAB). In the low probability context, the standard stimulus A is instead presented as a rare deviant (e.g., BBBBBABBBBBABBB). Comparisons of visual stimulus-evoked event-related potentials (ERPs) recorded using electroencephalography (EEG) reveal more negative-going waveforms evoked by deviants at posterior scalp electrodes from ~150-300 ms following stimulus onset (Czigler et al., 2004; Kimura et al., 2009; Stefanics et al., 2011). This difference in ERP waveforms is known as a visual mismatch response (VMR), or the visual mismatch negativity (vMMN; for recent reviews see Kimura, et al., 2011; Stefanics et al., 2014; Kremlacek et al., 2016). The vMMN is considered to be the visual counterpart of the earlier-discovered auditory MMN (Naatanen et al., 1978). VMRs are broadly theorised to reflect the tracking of patterns and regularities within sensory environments as implemented within the architecture of the visual system (e.g., Kimura et al., 2009).

Multiple phenomena can contribute to VMRs as they are measured using ERPs, each of which relates to different statistical regularities that are present in oddball sequences. The first class of phenomena relates to recent stimulation history, that is, which stimuli have recently been presented to an observer. Effects of immediate stimulus repetition, known as repetition suppression or stimulus-specific adaptation (Desimone, 1996; Movshon & Lennie, 1979), are defined as a reduction in a measure of neural activity (e.g., firing rates, local field potential amplitudes or BOLD signals) following repeated as compared to unrepeated stimuli (Grill-Spector, Henson & Martin, 2006). Repetition effects can partly account for VMRs as a suppression of neural responses evoked by standard stimuli and a lack of such suppression for the deviant stimuli (May & Tiitinen, 2010; Nelken & Ulanovsky, 2007). More recently developed excitatory-inhibitory circuit models also describe enhancement of responses to deviant stimuli that occur when both excitatory and lateral inhibitory responses are reduced following repeated exposure to standards (Dhruv, et al., 2011; Solomon & Kohn, 2014; Kaliukhovich & Vogels, 2016), where effects of disinhibition can occur over later time windows during the stimulus-evoked response than suppressive effects of stimulus repetition (Patterson et al., 2013). There are also systematic effects that depend on the combinations of stimuli that were recently presented to an observer (e.g., the most recently seen 2-5 stimuli, Sawamura et al., 2006; Vinken & Vogels, 2017; for similar effects on auditory evoked responses see Ulanovsky et al., 2004; Maheu et al., 2019).

The second class of phenomena relates to internally generated models of sensory environments that are learned through exposure to regularities in oddball sequences. In oddball sequences, expectations derived from these models are likely to be biased toward the often-appearing standard rather than the rare deviant. Such expectations are fulfilled when the standard is presented, and violated when a deviant appears in the expected standard’s place (Friston, 2005; Stefanics et al., 2014, 2018). There is ample evidence from non-oddball designs that an observer’s expectations can influence responses of stimulus-selective visual neurons (Summerfield et al., 2008; Amado et al., 2016; Hall et al., 2018; Feuerriegel et al., 2018a), and that humans can generate neural representations of expected sensory events even before the corresponding stimulus has been presented (Kok et al., 2017; Blom et al., 2020).

Within this prediction-based framework, there are two ways in which an observer’s learned expectations might relate to the standard/deviant ERP differences indexed by VMRs. The first account describes VMRs as a marker of prediction error, indexing the degree of mismatch between the predictions of an observer’s internal model and the actual sensory input (Friston et al., 2005; Garrido et al., 2009; Stefanics et al., 2014, 2018). VMRs are conceptualised as a combination of suppressed neural responses for subjectively expected standard stimuli and enhanced responses to surprising deviant stimuli. Variants of this model also incorporate the notion of precision (Feldman & Friston, 2010), whereby larger differences in the proportions of standard/deviant stimuli (and stronger weighted expectations to see standards) lead to larger prediction errors for deviant stimuli, and larger VMRs.

An alternative view is that VMRs index a surprise response that occurs when expectations have been violated, but not concurrent suppression when expectations are fulfilled. This account is equivalent to models which describe mismatch responses that occur following violations of regularities and recurring patterns in oddball sequences (e.g., Winkler, 2007; Paavilainen et al., 2013). Similar surprise responses also follow novel, unexpected or potentially rewarding stimuli in reward-learning paradigms, and precede the computation of reward prediction errors (reviewed in Schultz, 2016).

It has been remarkably difficult to isolate and quantify effects of immediate stimulus repetition, fulfilled expectations, and surprise using visual oddball sequences. This is because in such designs, the expected stimulus (the standard) is almost always also a repeated stimulus, whereas the surprising deviants are seldom repeated. In addition, repetition and expectation effects are likely to interact, whereby differences between expected and surprising stimuli are absent or diminished for immediately repeated stimuli (e.g., Todorovic & de Lange, 2012; Wacongne et al., 2011; Feuerriegel et al., 2018b). Recently-developed designs have attempted to overcome these limitations by adding sequences in which many different stimulus images (including the deviants and standards) are randomly interspersed within a sequence, known as equiprobable control sequences (e.g., Jacobsen & Schroger, 2001; Amado & Kovacs, 2016; reviewed in Stefanics et al., 2014). In equiprobable sequences, each stimulus image is presented with the same probability as the deviant stimuli in the classical oddball sequences. Expectations cannot be reliably formed for any particular stimulus image in the sequence, so the stimuli are neither expected nor surprising. ERPs evoked by the same stimulus image are then compared across equiprobable and deviant contexts to isolate surprise effects while apparently controlling for immediate repetition effects.

These studies have reported separable contributions of repetition suppression and surprise to VMRs (Amado & Kovacs, 2016; Astikainen, et al., 2008; Czigler et al., 2002; Kimura et al., 2009). However, these designs do not adequately isolate effects of fulfilled expectations from repetition effects, as the expected standard stimuli are repeated frequently whereas the equiprobable control stimuli are not. There are also two additional potential confounds to consider when using this design. Massed repetition of standards is also likely to enhance responses to deviants through reductions in lateral inhibition (Dhruv et al., 2011; Kaliukhovich & Vogels, 2016), and this response enhancement would not systematically occur in equiprobable sequences. The number of stimulus identities is also different across classical oddball sequences (2 identities) and equiprobable sequences (typically ~10 identities). This can produce systematic effects on visual evoked ERPs during the same time range as VMRs (Feuerriegel et al., 2018a), which have been localised to ventral temporal areas of the visual cortex (Pajani et al., 2017; Rostalski et al., 2020). It is unclear to what extent these confounds have contributed to existing reports of VMRs identified using equiprobable sequences. This has made it extremely difficult to cleanly isolate and quantify each repetition and expectation effect, which is critical for constraining theoretical models of VMRs.

To overcome these limitations, we recently developed an oddball design based on Fast Periodic Visual Stimulation (FPVS; Dzhelyova & Rossion, 2014a, 2014b; Dzhelyova, et al., 2017; Liu-Shuang, et al., 2014, 2016) that allowed us to isolate and quantify [surprising - expected] ERP differences while controlling for immediate stimulus repetition effects (Feuerriegel et al., 2018b). In FVPS oddball designs, a base stimulus is presented at a rapid, periodic rate (e.g., 6 Hz). Oddball stimuli replace the base stimulus every *N* stimuli at a fixed periodicity (e.g., every 7 stimuli). Viewers exposed to these stimulation sequences can perfectly predict when an oddball stimulus will appear (e.g., after 6 base rate stimuli have been presented). In sequences in which there are two possible oddball stimulus images, with one image appearing more often than the other, observers can also form expectations about the specific identity of each upcoming oddball stimulus. This leads to oddball stimulus-evoked VMRs that appear with similar topographies and latencies to VMRs found when using equiprobable sequences (Feuerriegel et al., 2018b). As six base face images are presented between each oddball face, the visual system is strongly adapted to the base face image before the presentation of each oddball stimulus in the sequences. This means that the state of adaptation in the visual system is equivalent for expected and surprising oddballs, thereby controlling for effects of immediate repetition and recent stimulation history.

In our previous work we identified [surprising – expected] response differences that were independent of stimulus repetition effects, however we could not isolate effects of surprise from those of fulfilled expectations. In the current study, we presented sets of FPVS oddball sequences and systematically varied the relative proportions of different oddball stimulus images in each sequence type, ranging from 10% to 90%. Critically, this included a 50% ‘expectation neutral’ condition, in which participants would not form strong expectations about which specific oddball stimulus was most likely to appear, as each oddball image appeared with equal probability. To isolate and quantify effects of fulfilled expectations and surprise, we compared ERPs evoked by expected (60-90% probability) and surprising (10-40% probability) oddballs with those evoked by neutral (50% probability) oddballs. We could also test whether neural responses indexed by VMRs varied in accordance with the *degree* of expectedness or surprise associated with seeing a given oddball stimulus (i.e. the objective oddball stimulus presentation probability).

We expected to find ERP differences for surprising compared to neutral conditions, consistent with surprise effects found in studies using equiprobable sequences (e.g., Kimura et al., 2009; Amado et al., 2016). We did not make explicit predictions regarding the magnitude of expectation suppression effects; however, we expected these to be of smaller magnitude than surprise effects, based on existing work using non-oddball designs (e.g., Amado et al., 2016). We also hypothesised that we would find linearly more negative-going ERP amplitudes for oddballs with lower presentation probabilities (i.e. more negative amplitudes for oddballs which were less probable and more surprising), consistent with the notion of precision-weighted prediction error signalling (Stefanics et al., 2014) and previous observations of graded effects on N2 and P3 ERP component amplitudes in oddball designs (e.g., Duncan-Johnson & Donchin, 1977; Polich & Margala, 1997; Wessel & Huber, 2019).

By controlling for immediate repetition effects, we could also test for adaptation effects that might occur over longer timescales, which may be important to consider in future experimental and computational modelling work. We assessed whether oddballUevoked neural response amplitudes gradually changed over the course of a stimulation sequence, as previously reported when using both classical auditory oddball (Ulanovsky et al., 2004) and FPVS oddball designs (Feuerriegel et al., 2018b).

## 2. Method

### 2.1. Participants

We report how we determined our sample size, all data exclusions (if any), all data inclusion/exclusion criteria, whether inclusion/exclusion criteria were established prior to data analysis, all manipulations, and all measures in the study.

Forty-eight people participated in this experiment. We originally planned to recruit 50 participants so that we would retain at least 40 datasets after exclusion of excessively noisy EEG data. We did not collect the final two datasets due to budget constraints. We aimed to recruit a similar sample size to our previous study, which reported expectation effects in the absence of immediate stimulus repetition confounds (Feuerriegel et al., 2018a). Participants were fluent in spoken and written English and had normal or corrected-to-normal vision. Five participants were excluded due to excessively noisy EEG data (i.e. more than 10 electrodes with a high number of large-amplitude artefacts). Forty-three participants were included in the final sample (20 males, age range 18-38 years, mean age 24.4 years, two left-handed). Participants were reimbursed 35 AUD for their participation in the experiment. The study was approved by the Human Research Ethics Committee of the Melbourne School of Psychological Sciences (Ethics ID 1953637).

### 2.2. Code and Data Availability

Code and data used for running the experiment and performing the analyses described here will be made available at https://osf.io/x8wfv/ at the time of publication.

### 2.3. Stimuli

Fifteen frontal images of faces (six male, nine female, neutral expression) were taken from the stimulus set in Laguesse and Rossion (2013). We converted the images to greyscale and equated their mean pixel intensity and RMS contrast using the SHINE toolbox (Willenbockel et al., 2010). Stimuli subtended approximately 5.4° × 7.9° visual angle at a viewing distance of 80 cm. We also scaled the resulting face images by 120% to create a separate set of images, which were used as oddball stimuli.

Faces are an ideal stimulus type for use in oddball designs, as they are associated with robust face identity repetition and expectation effects (Summerfield et al., 2008; Grotheer & Kovacs, 2015; Henson, 2016; Feuerriegel et al., 2018b). Moreover, in FPVS oddball designs, faces elicit a complex EEG response to changes of identity which has been well characterised in previous work (Dzhelyova & Rossion, 2014a, 2014b).

### 2.4. Procedure

Study methods and analyses were not pre-registered prior to the research being conducted. Participants sat in a dark room 80 cm in front of a gamma-corrected 24-inch LCD monitor (refresh rate 60 Hz). Stimuli were presented against a grey background matched to the mean pixel intensity of the images. A blue fixation cross was superimposed over the nasion of the face images throughout each sequence. Stimuli were presented using Psychtoolbox (Brainard, 1997; Kleiner et al., 2007) and MATLAB (The Mathworks, Natick MA, USA).

Each stimulation sequence began with the presentation of a fixation cross for two seconds, followed by an 86-second stimulation period, and then another two-second period during which only the fixation cross was visible. During the stimulation period, we presented face images at a rate of 6 Hz using square wave contrast modulation with a 50% duty cycle. At the beginning of each 166.66 ms image cycle, a face image appeared on the screen for 83.33 ms and was then replaced by the grey background for 83.33 ms. During the first two seconds of the stimulation period, the stimulus contrast gradually increased from 0 to 100% (i.e. a fade-in), and gradually decreased to 0% across the last two seconds of the stimulation period (i.e. a fade-out). Between each stimulation sequence participants could take a self-paced break (minimum possible break duration was 10 seconds).

In each sequence, a single identity base face appeared at a rate of 6 Hz (see Figure 1). Oddball faces of the same gender replaced the base face image for every 7^th^ image in the sequence. Oddball faces were always a different identity to the base face and 20% larger in size. Within each sequence, the oddball stimulus could be one of two face identities. The relative presentation proportions of each oddball face identity were manipulated across five sequence types, with expected/surprising oddball appearance probabilities of 90%/10%, 80%/20%, 70%/30%, 60%/40% and 50%/50%. For example, in the 90%/10% sequences one face identity would appear as 90% of oddballs, and the other face identity would be presented as 10% of oddballs. In these sequences, we anticipated that participants would form expectations to see the more common oddball face identity. By contrast, in the 50%/50% sequences, we did not expect participants to form expectations for a given oddball face, as both identities appeared with equal probability. We termed oddball images that were more than 50% to appear as expected (60-90% conditions), those less than 50% likely to appear as surprising (10-40% conditions), and oddballs with 50% appearance probability as neutral.

**Figure 1.**
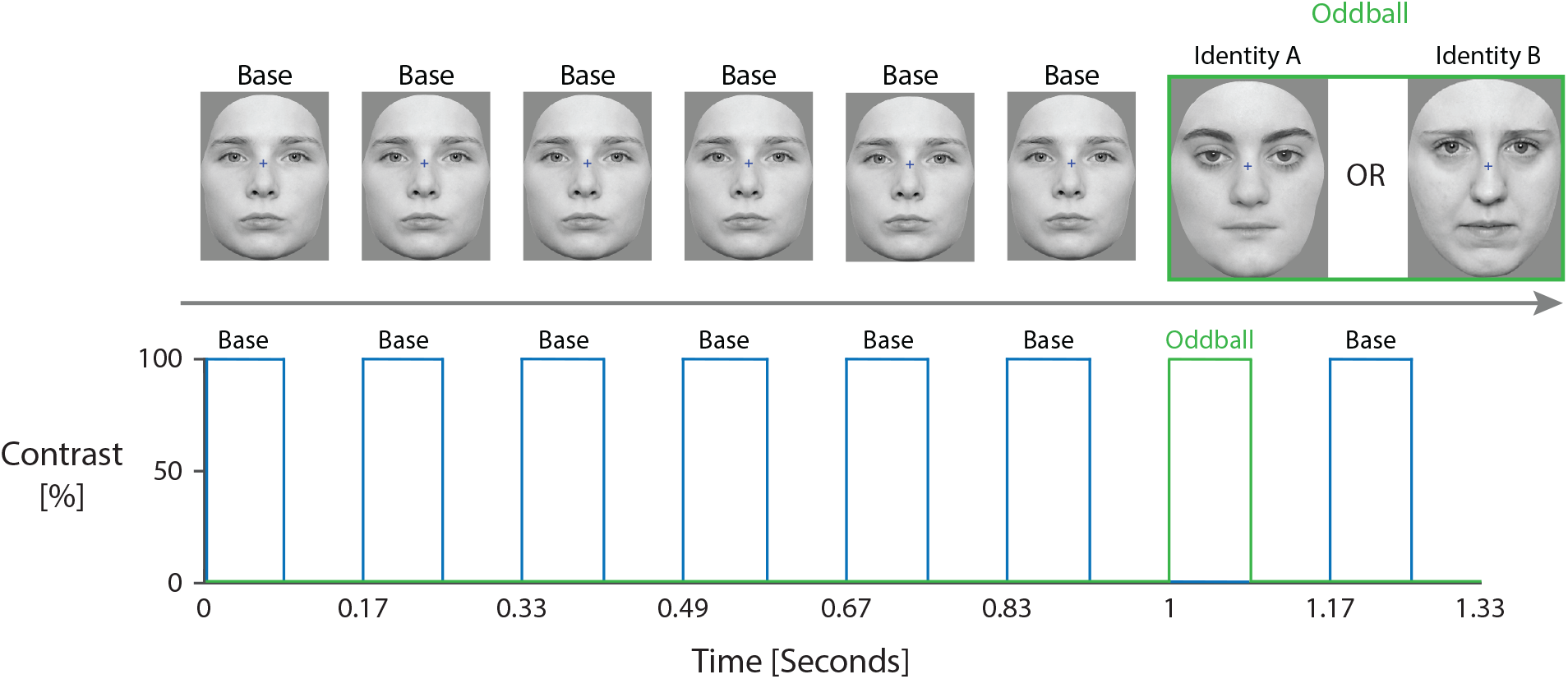
Stimulation sequence diagram. Faces were presented using square wave contrast modulation at a rate of 6 cycles per second (i.e. 6 Hz). After every six base faces, participants saw one of two possible oddball face identities. The presentation probabilities of each oddball identity varied from 10-90% in steps of 10% across sequences (e.g., one sequence type had 10% identity A, 90% identity B). This included an ‘expectation neutral’ sequence type in which both oddball identities were presented as oddballs with 50% probability.

To aid learning of the relative oddball probabilities, sequences were blocked by sequence type. Blocks of 6 consecutive sequences were presented for all sequence types except the 90/10 sequence type, which comprised a block of 10 sequences to ensure an adequate number of 10% probability oddballs per participant. To avoid abrupt transitions in the extent of the common/rare oddball probability splits across sequence types (e.g., a transition from a 90%/10% sequence to a 50%/50% sequence) half of the participants were first presented with a block of the 90%/10% sequence type, followed by blocks of 80%/20%, 70%/30%, 60%/40% and then 50%/50% sequence types. Block order was reversed for the other half of participants, starting with 50%/50% sequences and ending with 90%/10% sequences. Different sets of faces (one base face identity and two oddball identities) were presented as stimuli for each of the five sequence types. Different faces were used across sequence types so that participants expectations to see a specific face identity as the oddball were not transferred across successively presented sequence types. The sets of base/oddball faces allocated to each sequence type were counterbalanced across participants.

A total of 34 sequences of 90 seconds in duration were presented during the experiment. Total testing duration was 59 minutes excluding EEG equipment setup and self-paced breaks.

### 2.5. Experiment Task

We used a task that was orthogonal to the face stimuli in order to engage participants attention throughout the experiment (as done by Rossion & Boremanse, 2011; Liu-Shuang et al., 2014). Participants were instructed to fixate on a blue cross overlaying the face images and press a key on a TESORO Tizona numpad (1,000 Hz polling rate) when the cross changed colour from blue to red (ten colour changes per sequence, 100 ms colour change duration, minimum of two seconds between colour changes). We considered key presses within 1,000 ms of a colour change as correct responses. Numbers of correctly identified targets and mean response times were presented as feedback at the end of each sequence.

### 2.6. Task Performance Analyses

We calculated mean proportion correct scores and mean response times for correctly detected targets separately for each sequence type. We tested for differences in response times across sequence types using a one-way repeated measures ANOVA with the factor of sequence type (90/10, 80/20, 70/30, 60/40 and 50/50) with Greenhouse-Geisser corrections to account for violations of the sphericity assumption.

### 2.7. EEG Acquisition and Data Processing

We recorded EEG from 64 active scalp electrodes using a Biosemi Active Two system (Biosemi, the Netherlands). Recordings were grounded using common mode sense and driven right leg electrodes (http://www.biosemi.com/faq/cms&drl.htm). We added six channels to the standard montage: two electrodes placed 1 cm from the outer canthi of each eye, two electrodes placed above and below the left eye, and two electrodes over the left and right mastoid processes. EEG was sampled at 512 Hz (DC-coupled with an anti-aliasing filter, −3 dB at 102 Hz). Electrode offsets were kept within ± 50 μV.

We processed EEG data using EEGLab v13.4.4b (Delorme & Makeig, 2004). We identified excessively noisy channels by visual inspection (mean noisy channels per participant was 1.6, median = 1, range 0-5) and we excluded these from average reference calculation and independent components analysis (ICA). We also marked and removed time windows of data which contained large amplitude artefacts. We then re-referenced the data to the average of all channels and removed one extra channel (AFz) to correct for the rank deficiency caused by the average reference (as done by Feuerriegel et al., 2018a). The resulting data were low-pass filtered at 30 Hz (EEGLab Basic Finite Impulse Response Filter New, non-causal zero-phase, −6 dB cutoff frequency 33.75 Hz, transition bandwidth 7.5 Hz). We processed a copy of this dataset in the same way, but additionally applied a 0.1 Hz high-pass filter (same filter type as above, −6 dB cutoff frequency 0.05 Hz, transition bandwidth 0.1 Hz) to improve stationarity for the ICA. ICA was performed on this 0.1 Hz high-pass filtered dataset (RunICA extended algorithm, Jung et al., 2000). We then copied the independent component information to the dataset that had not been high-pass filtered. We identified and removed independent components generated by blinks and saccades according to guidelines in Chaumon et al (2015). After subtracting these components, we high-pass filtered the data at 0.1 Hz and interpolated any bad channels and AFz using the cleaned data (spherical spline interpolation).

The resulting EEG data were segmented from −166 ms to 1,000 ms relative to the onset of each oddball face, and baseline-corrected using the average −166 to 0 ms relative to oddball onset. This baseline window captured the period of one complete stimulus on/off cycle. Epochs containing amplitudes at any scalp electrode that exceeded ±100 μV were removed. Mean numbers of retained epochs per participant ranged from 67 (for 10% probability oddballs) to 606 (for 90% oddballs). Summary statistics related to numbers of epochs per condition are displayed in Supplementary Table S1.

### 2.8. Identification of VMR Regions of Interest

We first identified electrodes and time windows that would comprise our regions of interest (ROIs). To do this, we averaged oddball-evoked ERPs across epochs within each participant and each stimulus appearance probability condition. We then averaged the resulting ERPs across surprising (10-40% probability) and expected (60-90% probability) oddball types separately. We compared ERPs following expected and surprising oddballs at each time point for each of the 64 scalp electrodes using paired-samples t tests (alpha level 0.05). From these maps of significance-thresholded differences we visually identified groups of electrodes at which there were differences that persisted over an extended time window (i.e., at least 50 ms) to exclude isolated effects at single time points, which are likely to be false positives. Channel-by-timepoint matrices of statistically significant [surprising – expected] differences are displayed in Supplementary Figure S2.

At the ROI definition step, we did not correct for multiple comparisons. Here, we aimed to maximise our ability to identify any time windows over which VMRs may occur. Effects of stimulus-specific expectations tend to be small when controlling for immediate stimulus repetition effects (e.g., Feuerriegel et al., 2018a), and highly conservative correction methods may have prevented us from identifying any ROIs for subsequent analyses despite our substantial (n=43) dataset. Cluster-based permutation tests (e.g., Maris & Oostenveld, 2007), which retain higher sensitivity to detect real effects, are also not ideal for detecting visual mismatch effects that occur over a small number of parieto-occipital electrodes (e.g., the visual mismatch effect in Feuerriegel et al., 2018b). This is because these electrodes lie on the edge of the EEG cap and have few spatial neighbours (see Groppe et al., 2011). We acknowledge that this increases that chance of one of more ROIs reflecting false positive effects. However, we note that the visual mismatch effect corresponding to our earlier (200-350 ms) ROI was also identified in our previous work using the same face set and a highly similar experiment design (Feuerriegel et al., 2018b).

Subsequent analyses were based on the average ERP amplitudes across channels and time points within each identified ROI. To avoid inflation of false positive rates associated with circularity (see Kriegeskorte et al., 2009) all further analyses of ROI-averaged amplitudes did not involve direct comparisons across surprising and expected conditions. Each ROI-averaged mean amplitude analysis involved a comparison of either an expected or surprising oddball condition with the 50% expectation neutral condition (which was not included in the ROI selection step). All regression models described below were separately fit to data within surprising, neutral and expected conditions.

### 2.9. ERP Mean Amplitude Analyses

To separately quantify effects of expectation suppression and surprise, we used frequentist and Bayesian paired-samples t-tests (two-tailed) using JASP v0.9.1 (JASP Core Team) to estimate ERP amplitude differences between expected and surprising oddballs compared to neutral (50% probability) oddballs. Bayesian t-tests used a Cauchy distribution with width 0.75 (default settings in JASP). We then compared amplitudes for each individual oddball appearance probability condition to the neutral condition using paired-samples t tests.

As we had previously found linear changes in ERP amplitudes over the course of long-duration (e.g., 90-second) FPVS stimulation sequences (Feuerriegel et al., 2018b), we tested for linear relationships between oddball position in the sequence (ranging from the first oddball to the 70^th^ oddball) and ROI-averaged mean amplitudes. We then performed frequentist and Bayesian one-sample t tests to assess whether the resulting beta values (i.e. the slopes) corresponding to the effect of oddball position differed from zero. In these analyses, beta values larger than zero indicate more positive-going ERP amplitudes for oddballs presented later in a sequence. As we identified systematic effects of oddball position, this predictor was included in the regression analyses described below.

We additionally tested whether VMR amplitudes vary by objective oddball presentation probability, to assess whether VMRs signal information about the *degree* of expectedness or surprise associated with seeing a particular stimulus. We tested for linear relationships between stimulus appearance probability and ROI-averaged ERP amplitudes for surprising (10-40% probability) oddballs. This was not done for expected oddballs, as there was little evidence of expectation suppression effects in the data. To do this, we fit linear regression models to patterns of single-epoch ROI-averaged amplitudes separately for each participant dataset, using predictors of stimulus appearance probability (ranging from 10-40%) and oddball position in the sequence (ranging from 1-70). We then tested whether beta values differed from zero at the group level using frequentist and Bayesian single-sample t tests as described above. In these analyses, beta values larger than zero indicate more positive-going amplitudes for oddballs with higher presentation probabilities.

## 3. Results

### 3.1. Task Performance

Target detection rates for the fixation cross colour change detection task were near ceiling for most participants. Group mean proportion correct scores ranged between 95-97% across sequence types (individual participant data displayed in Supplementary Figure S1A). Group mean RTs for correctly detected targets were also very similar and ranged between 372-381 ms across sequence types (Supplementary Figure S1B). There was no statistically significant effect of sequence type on mean RTs, F(2.6, 107.6) = 2.46, p = 0.076. Overall, these results do not indicate any substantial differences in task performance across sequence types.

### 3.2. Identification of VMR Regions of Interest

We visually identified two time windows during which there were more negative-going amplitudes following surprising compared to expected oddballs: an early time window (200-350 ms) and a late time window (500-1,000 ms; group-averaged ERPs are depicted in Figure 2A). For both time windows, differences were identified at bilateral occipitoparietal electrodes P7, P8, P9, P1S, PO7 and PO8 (for scalp maps of time window-averaged mean amplitudes see Figure 2B). The latency and topography of the early effect closely resembles VMRs observed in our previous FPVS oddball experiment (Feuerriegel et al., 2018b), as well as VMRs reported when using equiprobable control sequences (Kimura et al., 2009; Amado & Kovacs, 2016). In addition, we also observed more positive-going ERPs for surprising stimuli at midline central and parietal electrodes for the early and late time windows (electrodes POz, PO3, Pz, P1, P2, P4, CP3, CPz, CP1, CP2, Cz, FCz, FC2). Analyses using these additional ROIs are presented in the Supplementary Material, and produced almost identical patterns of results to those found using the negative-going [surprising – expected] difference ROIs described above.

**Figure 2.**
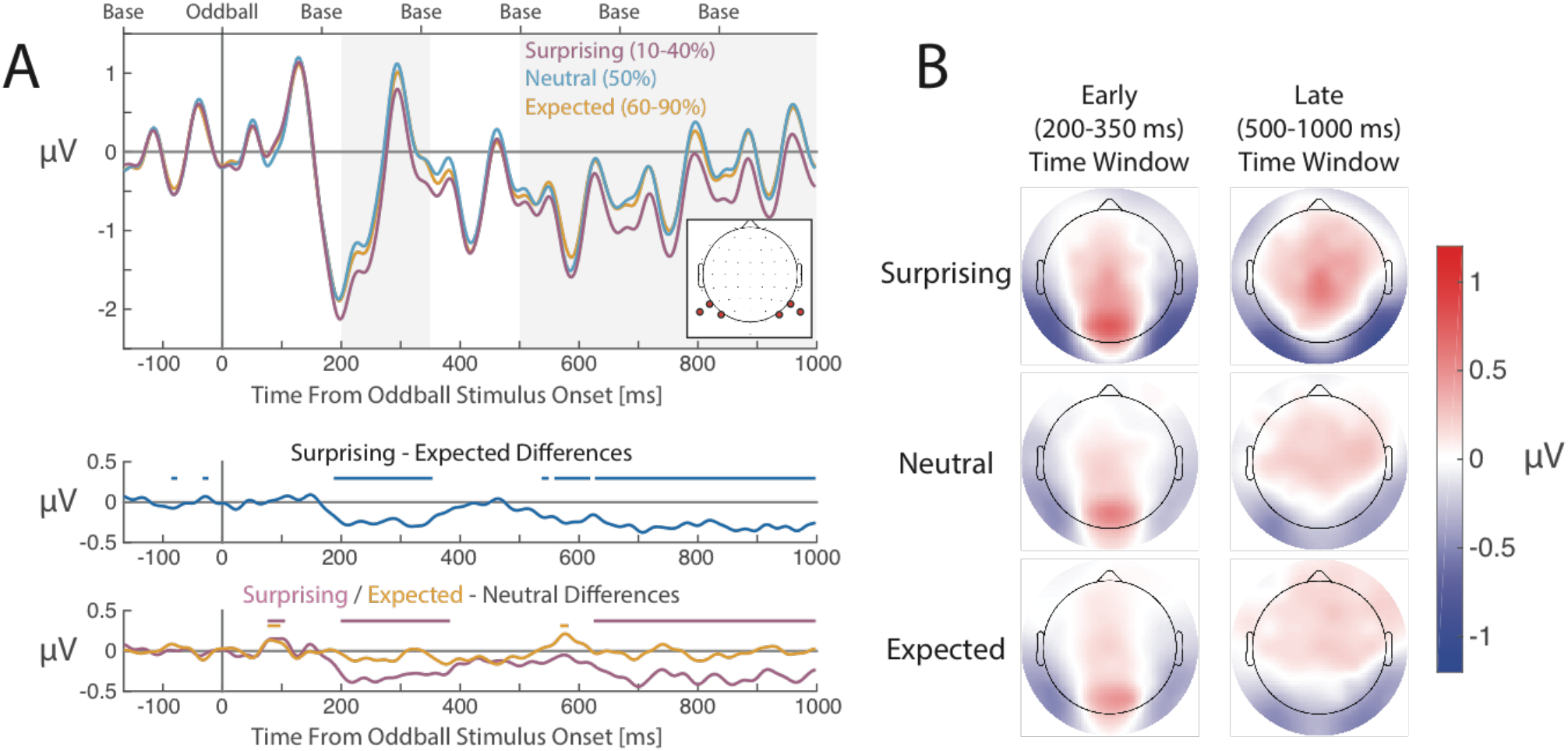
Group-averaged ERPs evoked by surprising (10-40% probability), neutral (50%) and expected (60-90%) oddballs. A) Group-averaged ERPs for each condition (upper panel), differences between surprising and expected conditions (middle panel), and differences between surprising and expected conditions compared to the neutral condition (lower panel), averaged across occipitopareital electrodes P7/8/9/10 and PO7/8 (electrode locations shown in the inset scalp map). Grey shaded areas denote the early (200-350 ms) and late (500-1000 ms) ROI time windows. Labels at the top of the upper panel denote the onset times of the base and oddball stimuli within the epoch. Thick lines at the top of each plot denote statistically significant differences between ERPs that were averaged over selected occipitoparietal electrodes (p < 0.05 uncorrected). Statistical significance markers for the [surprising - expected] contrast may have an inflated false positive rate as electrodes used for plotting ERPs were selected based on identification of significant [surprising - expected] differences. B) Scalp maps of group-averaged mean amplitudes for surprising, neutral and expected conditions, for the early (200-350 ms) and late (500-1,000 ms) time windows. Note that this figure depicts ERPs, difference waves and scalp maps calculated after removal of epochs that were preceded by a surprising oddball. Numerical differences between surprising and expected oddball conditions may be inflated due to the fact that both ROIs were defined based on significant [surprising - expected] condition differences.

Differences between surprising/neutral and expected/neutral conditions are plotted in the bottom panels of Figure 2A. There were clearly-visible differences between surprising and neutral conditions (i.e. surprise effects) during the early and late time windows, but not between expected and neutral conditions. As surprise effects lasted until the end of the 1,000 ms epoch, we were concerned that these effects would systematically influence pre-stimulus baselines for oddballs that follow a surprising oddball. We therefore excluded epochs corresponding to oddballs that were preceded by a surprising oddball (e.g., as done by Kimura et al., 2009). Numbers of useable epochs per condition remained high following epoch selection (see Supplementary Table S2). Note that the averaged ERP waveforms, difference waves and scalp maps displayed in Figure 2 were calculated after excluding epochs that were preceded by a surprising oddball.

As noted in section 2.8, these ROIs were defined based on results of paired-samples t tests that were not corrected for multiple comparisons. We also applied a false discovery rate control procedure (Benjamini & Hochberg, 1995) using the Mass Univariate ERP Toolbox (Groppe et al., 2011). Channel-by-timepoint matrices for [surprising - expected], [surprising - neutral] and [expected - neutral] contrasts are plotted in Supplementary Figure S2. In these plots there were negative-going [surprising - expected] differences at electrodes P9 and P10 the early ROI time window. Similar effects could be observed at electrode P10 during the late time window. The spatiotemporal pattern of [surprising - neutral] differences closely resembled the pattern of [surprising - expected] differences. No [expected - neutral] differences were statistically significant after controlling the false discovery rate.

### 3.3. Quantifying Effects of Fulfilled Expectations and Surprise

To isolate and quantify effects of expectation suppression and surprise, we compared mean amplitudes for expected and surprising oddballs with those for neutral oddballs. Mean amplitudes were more negative-going for surprising compared to neutral oddballs for the early time window (p = 0.005, BF_10_ = 6.78) and the late time window (p < 0.001, BF_10_ = 1141.24). By contrast, amplitudes did not appear to differ between expected and neutral conditions, with Bayes factors indicating evidence for the null hypothesis (early time window: p = 0.433, BF_10_ = 0.22, late time window: p = 0.966, BF_10_ = 0.17).

We next performed analyses at a more granular level by comparing the neutral condition with each other stimulus appearance probability condition, to assess whether this pattern of results was consistent with the condition-averaged data. Scalp maps of mean amplitudes by stimulus appearance probability condition, averaged over early and late time windows for each condition, are displayed in Figure 3A. Scatterplots of mean amplitudes by stimulus probability are displayed in Figures 3B (early time window) and 3C (late time window). Results of paired-samples t tests are displayed in Supplementary Table S3 for the early time window, and Supplementary Table S4 for the late time window. ROI-averaged amplitudes were more negative-going for all surprising oddball conditions across both time windows (p’s < 0.05) except for the 20% appearance probability condition (early time window p = 0.092, late time window p = 0.062). Group mean [surprising - neutral] difference magnitudes were similar across all surprising oddball conditions, ranging from −0.26 to −0.39 μV (early time window) and −0.20 to −0.40 μV (late time window). By contrast, none of the expected oddball conditions evoked significantly more positive or negative amplitudes compared to the neutral condition (all p’s > 0.258) except the 60% appearance probability oddballs, which evoked more negative-going amplitudes during the early time window (p = 0.035). Group mean [expected - neutral] difference magnitudes were much smaller than the observed surprise effects, and ranged from −0.21 to 0.08 μV (early time window) and −0.03 to 0.02 μV (late time window).

**Figure 3.**
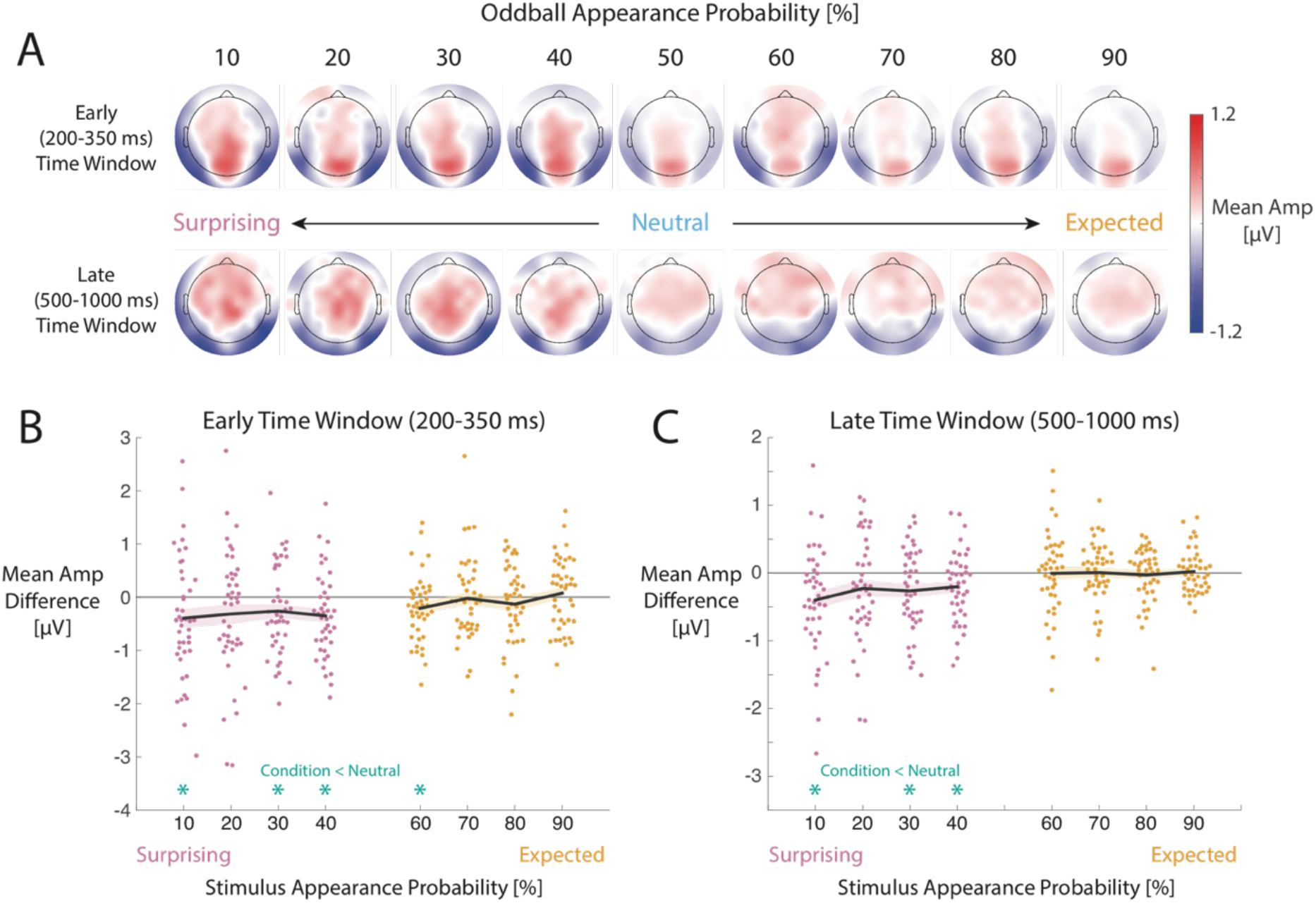
Mean amplitudes for early (200-350 ms) and late (500-1,000 ms) time windows, sorted by stimulus appearance probability. A) Scalp maps of mean amplitudes by stimulus appearance probability condition. B) Mean amplitude differences between each oddball appearance probability condition and the neutral (50%) condition. Thick black lines represent the group mean differences. Shaded regions represent standard errors. Asterisks at the bottom of each plot indicate statistically significant (p < 0.05) differences.

### 3.4. Graded Effects of Stimulus Appearance Probability

After identifying contributions of surprise to VMR amplitudes, we then assessed whether there were graded differences in ROI-averaged amplitudes by oddball presentation probability for surprising (10-40%) oddballs. In other words, we tested whether ERP amplitudes indexed an observer’s *degree* of surprise, which should differ across the 40% (slightly unlikely) and 10% (very unlikely) oddball types. For the early time window beta values did not significantly differ from zero, and the Bayes factor indicated moderate evidence for the null hypothesis, t(42) = 0.20, p = 0.847, BF_10_ = 0.17, mean beta value 0.001, CI [−0.01, 0.01]. Here, the mean estimated beta value denotes an average change of only 0.001 μV for each percentage increase in oddball appearance probability. We found similar results for the late time window, t(42) = 1.24, p = 0.221, BF_10_ = 0.34, mean beta value 0.005, CI [−0.003, 0.01].

### 3.5. Effects of Oddball Position in a Sequence

We also assessed whether there were systematic effects of the oddball position in the sequence (ranging from 1-70) on ROI-averaged amplitudes, as previously identified when using an FPVS oddball design (Feuerriegel et al., 2018b). For these analyses, we included all oddball presentations and did not exclude epochs that were preceded by a surprising oddball presentation.

For the early time window, beta values for the linear effect of oddball position were significantly above zero, t(42) = 10.19, p < 0.001, BF_10_ > 10,000, mean beta value 0.01, CI [0.009, 0.013], indicating more positive-going amplitudes for oddballs presented later in the sequence (depicted in Figure 4A) as reported in Feuerriegel et al. (2018b). Effects of oddball position also appeared be highly similar for expected, neutral and surprising oddball types (Figure 4B).

**Figure 4.**
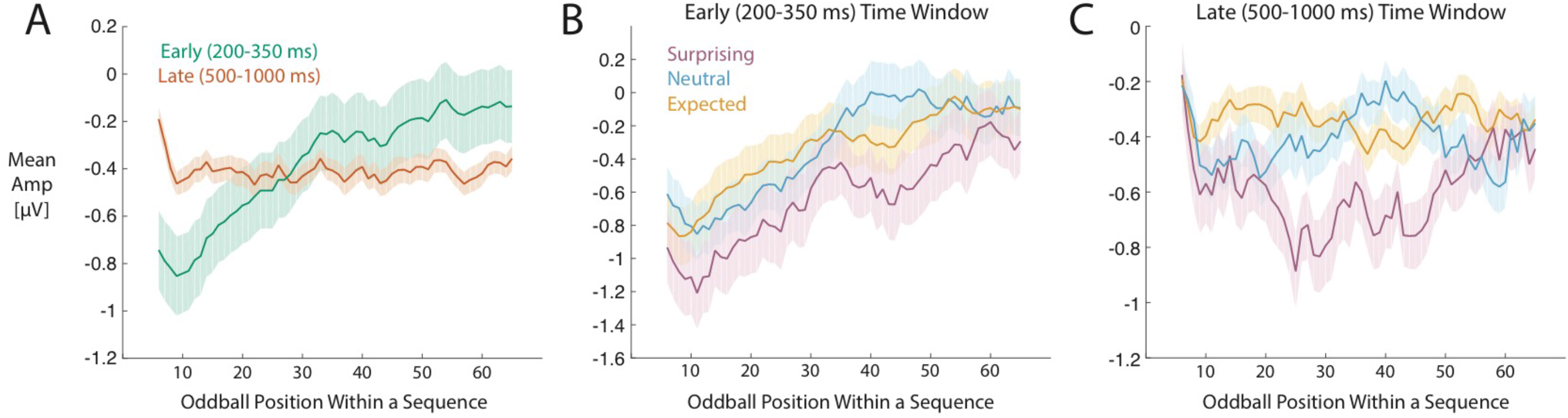
Group mean amplitudes by oddball position within each sequence, for early (200-350 ms) and late (500-1,000 ms) time window ROIs. A) Mean amplitudes by sequence, averaged across all stimulus appearance probability conditions. For the early time window, there is a clearly-visible linear trend toward more positive-going amplitudes for oddballs presented later in the sequences. Amplitudes are also slightly more negative-going for later oddballs during the late time window. B) Mean amplitudes for the early time window, plotted separately for surprising, neutral and expected conditions. C) Mean amplitudes for the late time window, plotted separately for surprising, neutral and expected conditions. For both time windows, effects of sequence position appear to be qualitatively similar for expected, neutral and surprising oddball types. Data has been smoothed with a boxcar function using the ten surrounding oddball positions. Shaded regions represent standard errors.

For the late time window, beta values were significantly below zero, t(42) = −3.16, p = 0.003, BF_10_ = 11.52, mean beta value −0.002, CI [−0.004, −0.001], and the effect was much smaller in magnitude than for the early time window (estimated average change of −0.002 μV for each unit increase in oddball position in the sequences). There were no clearly-visible differences in oddball position effects across surprising, neutral and expected conditions (Figure 4C).

## 4. Discussion

Using classical oddball designs it has been remarkably difficult to disentangle effects of immediate stimulus repetition, expectation and surprise in relation to visual mismatch responses (VMRs). By instead using a modified oddball design that controls for stimulus repetition effects, we could successfully isolate and quantify the contributions of expectation suppression and surprise. We report that VMRs as measured in visual oddball designs index a surprise response that occurs when an observer’ s expectations are violated. However, we did not find evidence of expectation suppression effects, in that oddball stimuli with very high probabilities of appearing (e.g., 90%) evoked ERP responses that were no different to an expectation neutral condition (Bayes factors indicated that any such effects were either absent or very small in magnitude). In addition, we did not find systematic variations in the extent of surprise responses by stimulus appearance probability (i.e. the likelihood of a certain stimulus appearing). In other words, VMRs were of similar magnitude for 40% probability (slightly unlikely) and 10% probability (very unlikely) oddballs. We also identified gradual changes in ERP amplitudes according to whether oddballs were presented relatively early or late within a stimulation sequence, as reported in previous work (Ulanovsky et al., 2004; Feuerriegel et al., 2018b). Our findings indicate that VMRs signal the occurrence of surprising or novel events, but convey little information about the *degree of* surprise associated with an event.

### 4.1. VMRs Appear to Index a Surprise Response, but Not Expectation Suppression

We found clear evidence for surprise effects; however, we did not observe expectation suppression. That is, ERP amplitudes were markedly different for surprising compared to neutral oddballs, but no differences were found between expected and neutral oddballs. This indicates that VMRs predominantly index a surprise response, and the relative magnitude of any effects of fulfilled expectations on ERP amplitudes (if they exist) are likely to be very small. These results also favour theoretical accounts that describe VMRs as signifying expectation violations (e.g., Winkler, 2007; Paavilainen et al., 2013), rather than predictive coding-based models which describe VMRs as reflecting both enhanced prediction error responses for deviants and suppressed prediction errors for expected standards (e.g., Friston, 2005; Stefanics et al., 2014, 2018). We believe that our findings extend to VMRs that share broadly similar scalp topographies and response latencies, which have been observed in previous work that used strategies to separate repetition effects from surprise responses (e.g., Astikainen et al., 2008; Kimura et al., 2009; Amado & Kovacs, 2016; File & Czigler, 2019; Feuerriegel et al., 2018b; reviewed in Kremlacek et al., 2016).

Based on our findings, we propose that VMRs in classical oddball sequences reflect contributions of immediate stimulus repetition and recent stimulus exposure (e.g., Ulanovsky et al., 2004; Sawamura et al., 2006; Maheu et al., 2019) that partially overlap in time with expectation violation responses that occur following deviant stimuli (e.g., Kimura et al., 2009; Amado & Kovacs, 2016). Response enhancement for deviants may also result from disinhibition of stimulus-selective visual neurons caused by adaptation to the often-repeated standard stimuli (e.g., Dhruv et al., 2011; Kaliukhovich & Vogels, 2016). Although we separated these effects in our experiment, they are likely to interact in the context of classical oddball sequences. Surprise responses appear to scale with the magnitude of stimulus-evoked responses, which can be suppressed via immediate stimulus repetition (Feuerriegel et al., 2018b) or changing the focus of spatial and feature-based attention away from the critical stimuli (File & Czigler, 2019; Smout et al., 2020). Here, we distinguish between the focus of spatial or feature-based attention and the task-relevance of the stimuli that evoke VMRs, as the critical stimuli in visual oddball designs are often not directly relevant to an experimental task.

Based on this evidence, the surprise responses indexed by VMRs appear to signal the presence of unexpected or novel *changes* in an observer’ s visual environment that are within the focus of attention. VMRs may preferentially signal salient events that are relevant for additional computations associated with statistical learning (Leshinskaya & Thompson-Schill, 2020), detection of potential rewards (Schultz, 2016), belief updating (Bennet et al., 2015) and adaptive changes to decisionmaking strategies following unexpected events (Wessel et al., 2017). VMRs might even index a process that enables or facilitates these different types of computations, similar to the proposed role of other types of surprise responses that precede computations of expected reward and subjective utility (discussed in Winkler, 2007; Schultz, 2016).

Our findings also suggest that, in some contexts, the notion of expectation suppression (Summerfield et al., 2008; Todorovic & de Lange, 2012; Grotheer & Kovacs, 2015; Pajani et al., 2017) may be a misnomer when used to explain neural response differences between expected and surprising stimulus contexts. Studies using non-oddball designs and probabilistic cueing have similarly reported much larger effects of surprise on fMRI BOLD signals than of fulfilled expectations (Egner et al., 2010; Amado et al., 2016; reviewed in Kovacs & Vogels, 2014). Previous studies using roving stimulus variants of oddball designs have reported expectation-related suppression of prediction error-related neural responses (e.g., Stefanics et al., 2018), however these designs tend to conflate expectation and repetition effects, and the observed neural response reductions could instead be attributed to repetition suppression. Where feasible, we recommend including expectation neutral conditions in future studies to quantify relative effects of expectation suppression and surprise.

### 4.2. Graded Effects of Stimulus Appearance Probability

Contrary to our hypotheses, surprise responses did not linearly vary with stimulus appearance probability. This indicates that VMRs as isolated in our experiment do not signal the *degree* to which a stimulus is surprising, but instead reflect an all-or-none fixed-amplitude response that occurs following expectation violations. This pattern does not fit with precision-based expectation suppression accounts of VMRs, which would predict more negative-going amplitudes for stimuli that are less likely to appear (e.g., Stefanics et al., 2018).

Our findings also contrast with previous reports of N2 and P3 ERP component amplitudes scaling with stimulus appearance probability (e.g., Duncan-Johnson & Donchin, 1977; Polich & Margala, 1997; Wessel & Huber, 2019). This is likely because the critical stimuli were task relevant in those studies, but not in our experiment. Task relevance of the critical stimuli may be an important factor that determines whether expectancy-related computations that occur following detection of surprising events (e.g., as specified in Schultz, 2016) are reflected in additional ERP components or other neural activity measures. These additional processes may exhibit a higher degree of sensitivity to differences in contextual event probability than those identified in our study.

### 4.3. Gradual Changes in ERP Amplitudes During Stimulation Sequences

In addition to the surprise effects described above, we also replicated findings of gradual changes in neural response amplitudes that occur over the course of a stimulation sequence (Ulanovsky et al., 2004; Feuerriegel et al., 2018b). This was indexed by more positive-going ERP amplitudes during the early VMR time window for oddballs presented later in the FPVS stimulation sequences, and slightly more negative-going amplitudes during the late VMR time window. These effects also appeared to be qualitatively similar for expected, neutral and surprising oddballs. The cause of these gradual amplitude changes over time is unclear, but may reflect the consequences of prolonged continuous sensory stimulation (discussed in Ulanovsky et al., 2004). As we had observed these effects in our previous work (Feuerriegel et al., 2018b), we distributed expected and surprising oddballs evenly throughout the stimulation sequences in order to avoid spurious findings that are due to this effect. Given that the magnitudes of these effects were large compared to the surprise effects in our study, we recommend that others test for such gradual changes over time and explicitly account for these in statistical and computational models, particularly when model parameters reflecting sequence learning also change over similar timescales.

### 4.4. Study Limitations

Our findings should be interpreted with the following caveats in mind. First of all, the timing and topography of the VMRs observed here may be particular to experiments that present faces as stimuli. We used faces as they tend to elicit robust repetition and expectation effects, and the time-course of face oddball-evoked ERPs has been well-characterised in previous work (e.g., Dzhelyova & Rossion, 2014a, Quek & Rossion, 2017). We have also previously used face stimuli to identify VMRs in an FPVS oddball design (Feuerriegel et al., 2018b) which has many hallmark features of VMRs observed when using other stimulus categories (e.g., Kimura et al., 2009; Astikainen et al., 2008; Amado & Kovacs, 2016). However, VMRs do appear to qualitatively differ depending on the stimulus features that are relevant to an observer’ s expectations (Amado & Kovacs, 2016; Stefanics et al., 2019; Robinson et al., 2020; Smout et al., 2020), possibly reflecting response amplitude changes in sensory neurons that are selective for those features. It remains to be seen whether visual mismatch responses that correspond to different anatomical locations within the visual hierarchy index different contributions of expectation suppression and surprise.

Although we controlled for immediate repetition effects, the effects of surprise observed in our study may index contributions of delayed repetition effects, which depend on the recency with which an observer had last seen a presented oddball face identity (e.g., Henson et al., 2004; De Baene & Vogels, 2010; Ulanovsky et al., 2004). For example, in higher presentation probability conditions it was more likely that the preceding set of oddball faces were of the same identity as the currently-presented oddball. It is unclear whether a slowly-decaying stimulus repetition effect (e.g., as modelled in Vinken & Vogels, 2017) would be sufficient to produce a step-like function between surprising (10-40%) and neutral (50%) conditions as observed in our study, rather than a monotonic effect of stimulus appearance probability. In any case, the potential influence of delayed repetition effects should not be ruled out when interpreting our findings.

Effects of recent stimulation history and stimulus expectations are difficult to disentangle in oddball designs because expectations are often entrained by manipulating the frequency of stimulus presentation. Notably, VMRs resembling those found in oddball designs (e.g., Kimura et al., 2009; Amado & Kovacs, 2016; Feuerriegel et al., 2018b) are not typically observed in probabilistic cueing designs that do not have systematic delayed repetition confounds (e.g., Summerfield et al., 2011; Feuerriegel et al., 2018a, Hall et al., 2018, but see Todorovic & de Lange, 2012 for such effects when presenting auditory stimuli). Direct comparisons of expectation-related effects across similar variants of each design may help to better specify those experimental manipulations that are responsible for VMRs.

In our design, six base rate faces were presented both before and after each oddball stimulus. This means that our results may also have captured effects of surprise that act on visual evoked responses to base faces presented after a surprising oddball. The effects over the early (200-350 ms) time window are similar to those identified in other designs that did not present additional stimuli immediately after a surprising face (e.g., Amado et al., 2016). However, it is plausible that effects observed in the later (500-1000 ms) time window could reflect altered visual evoked responses to base faces due to increased pupil dilation following surprising events (e.g., Richter & de Lange, 2019; reviewed in Wessel, 2017). More generally, the consequences of pupil dilation following surprising events also appears to substantially contribute to surprise effects on BOLD signals observed in non-oddball designs (Richter & de Lange, 2019).

In addition, we used a limited range of stimulus appearance probabilities ranging from 10% to 90%. Consequently, we cannot rule out graded effects of stimulus appearance probability that might be identified using sequences with more extreme expected/surprising probability splits. For example, Javitt et al. (1998) presented auditory deviants at probabilities ranging between 0.56% and 15%, and found progressively more negative-going mismatch response amplitudes in smaller deviant probability blocks (see also Pincze et al., 2002). Experiments presenting stimuli at very low probabilities should include a familiarisation period to avoid potential confounding effects of stimulus novelty that may differ between expected and surprising stimuli (reviewed in Schomaker & Meeter, 2015; Feuerriegel, 2016).

We also note that other phenomena than those identified here are likely to influence VMR magnitudes, and may operate over a broad range of timescales (see Ulanovsky et al., 2004; Sawamura et al., 2006; Maheu et al., 2019). Our openly-available dataset can be used to test for candidate effects, which can then be replicated in future work. The set of effects identified here can also be used to develop computational models of VMRs, which may account for complex sets of potentially interacting phenomena in a mathematically explicit fashion (e.g., Stefanics et al., 2018), however this is beyond the scope of the current study.

In our design there were also very different numbers of epochs across the stimulus appearance probability conditions, ranging from an average of 67 included epochs for 10% probability oddballs to an average of 531 epochs for 90% probability oddballs. The single time-point and ROI-averaged mean amplitude analyses reported here do not suffer from systematic biases as a function of the number of included epochs, in contrast to ERP component peak amplitude measures (Clayson et al., 2012). However, there was systematically larger across-participant variability in estimates for low compared to high probability oddball conditions (visible in the single participant data points in Figure 3B and 3C). These differences in across-subject variance could potentially lead to [expected - neutral] effects being easier to detect than [surprising - neutral] effects. We instead found the opposite pattern; there were robust [surprising - neutral] condition differences but no apparent [expected - neutral] differences in our data.

### 4.5. Conclusion

We report that VMRs index a surprise response that occurs when an observer’ s expectations about upcoming sensory events are violated. The magnitude of these surprise responses does not appear to vary with graded changes in stimulus appearance probability. Our findings suggest that VMRs may signal the presence of unexpected *changes* in the environment, functioning as a precursor to further computations that utilise surprise signals for statistical learning, reward seeking and adaptive changes to decision-making strategies. However, we did not find support for predictive coding accounts of VMRs that describe suppression of prediction error responses when an observer’s expectations are fulfilled.

## Acknowledgements

This project was supported by a University of Melbourne Early Career Researcher Grant awarded to D.F., a European Union Horizon 2020 Marie Sklodowska-Curie Individual Fellowship awarded to G.L.Q. (841909), and Australian Research Council (ARC) Discovery Project Grants awarded to H.H. (DP180102268) and S.B. (DP160103353). Funding sources had no role in study design, data collection, analysis or interpretation of results.

## References

Amado, C., & Kovacs, G. (2016). Does surprise enhancement or repetition suppression explain visual mismatch negativity? European Journal of Neuroscience, 43(12), 1590–1600.

Amado, C., Hermann, P., Kovács, P., Grotheer, M., Vidnyanszky, Z., & Kovács, G. (2016). The contribution of surprise to the prediction based modulation of fMRI responses. Neuropsychologia, 84, 105–112.

Astikainen, P., Lillstrang, E., & Ruusuvirta, T. (2008). Visual mismatch negativity for changes in orientation - A sensory memory-dependent response. European Journal of Neuroscience, 28(11), 2319–2324.

Benjamini, Y., & Hochberg, Y. (1995). Controlling the false discovery rate: A practical and powerful approach to multiple testing. Journal of the Royal Statistical Society Series B: Statistical Methodology, 57, 289–300.

Bennett, D., Murawski, C., & Bode, S. (2015). Single-trial event-related potential correlates of belief updating. eNeuro, 2(5), ENEURO.0076-15.2015.

Blom, T., Feuerriegel, D., Johnson, P., Bode, S., & Hogendoorn, H. (2020). Predictions drive neural representations of visual events ahead of incoming sensory information. Proceedings of the National Academy of Sciences of the U.S.A., 117(13), 7510–7515.

Brainard, D. H. (1997). The Psychophysics Toolbox. Spatial Vision, 10(4), 433–436.

Chaumon, M., Bishop, D. V., & Busch, N. A. (2015). A practical guide to the selection of independent components of the electroencephalogram for artifact correction. Journal of Neuroscience Methods, 250, 47–63.

Clayson, P.E., Baldwin, S.A., & Larson, M.J. (2012). How does noise affect amplitude and latency measurement of event-related potentials (ERPs)? A methodological critique and simulation study. Psychophysiology, 50, 174–186.

Czigler, I., Balazs, L., & Pato, L. G. (2004). Visual change detection: Event-related potentials are dependent on stimulus location in humans. Neuroscience Letters, 364(3), 149–153.

Czigler, I., Balazs, L., & Winkler, I. (2002). Memory-based detection of task-irrelevant visual changes. Psychophysiology, 39(6), 869–873.

De Baene, W., & Vogels, R. (2010). Effects of adaptation on the stimulus selectivity of macaque inferior temporal spiking activity and local field potentials. Cerebral Cortex, 20(9), 2145–2165.

Delorme, A., & Makeig, S. (2004). EEGLAB: An open source toolbox for analysis of single-trial EEG dynamics including independent component analysis. Journal of Neuroscience Methods, 134(1), 9–21.

Desimone, R. (1996). Neural mechanisms for visual memory and their role in attention. Proceedings of the National Academy of Sciences of the United States of America, 93(24), 13494–13499.

Dhruv, N.T., Tailby, C., Sokol, S.H., & Lennie, P. (2011). Multiple adaptable mechanisms early in the primate visual pathway. Journal of Neuroscience, 31(42), 15016–15025.

Duncan-Johnson, C.C., & Donchin, E. (1977). On quantifying surprise: The variation of event-related potentials with subjective probability. Psychophysiology, 15(5), 456–467.

Dzhelyova, M., & Rossion, B. (2014a). The effect of parametric stimulus size variation on individual face discrimination indexed by fast periodic visual stimulation. BMC Neuroscience, 15, 87.

Dzhelyova, M., & Rossion, B. (2014b). Supra-additive contribution of shape and surface information to individual face discrimination as revelaed by fast periodic visual stimulation. Journal of Vision, 14(14): 15. 1–14.

Dzhelyova, M., Jacques, C., & Rossion, B. (2017). At a single glance: Fast Periodic Visual Stimulation uncovers the spatio-temporal dynamics of brief facial expression changes in the human brain. Cerebral Cortex, 8(1), 4106–4123.

Egner, T., Monti, J. M., & Summerfield, C. (2010). Expectation and surprise determine neural population responses in the ventral visual stream. Journal of Neuroscience, 30(49), 16601–16608.

Feldman, H., & Friston, K. J. (2010). Attention, uncertainty, and free-energy. Frontiers in Human Neuroscience, 4, 215.

Feuerriegel, D. (2016). Selecting appropriate designs and comparison conditions in repetition paradigms. Cortex, 80, 196–205.

Feuerriegel, D., Churches, O., Coussens, S., & Keage, H.A.D. (2018a). Perceptual expectations do not modulate image repetition effects as measured by event-related potentials. Neuroimage, 169(1), 94–105.

Feuerriegel, D., Keage, H. A. D., Rossion, B., & Quek, G. L. (2018b). Immediate stimulus repetition abolishes stimulus expectation and surprise effects in fast periodic visual oddball designs. Biological Psychology, 138,110–125.

File, D. & Czigler, I. (2019). Automatic detection of violations of statistical regularities in the periphery is affected by the focus of spatial attention: A visual mismatch negativity study. European Journal of Neuroscience, 49(10), 1348–1356.

Friston, K. (2005). A theory of cortical responses. Philosophical Transactions of the Royal Society of London. Series B: Biological Sciences, 360(1456), 815–836.

Garrido, M. I., Kilner, J. M., Stephan, K. E., & Friston, K. J. (2009). The mismatch negativity: A review of underlying mechanisms. Clinical Neurophysiology, 120(3), 453–463.

Grill-Spector, K., Henson, R., & Martin, A. (2006). Repetition and the brain: Neural models of stimulus-specific effects. Trends in Cognitive Sciences, 10(1), 14–23.

Groppe, D. M., Urbach, T. P., & Kutas, M. (2011). Mass univariate analysis of event-related brain potentials/fields I: A critical tutorial review. Psychophysiology, 48, 1711–1725.

Grotheer, M., & Kovacs, G. (2015). The relationship between stimulus repetitions and fulfilled expectations. Neuropsychologia, 67, 175–182.

Hall, M.G., Mattingley, J.B., & Dux, P.E. (2018). Electrophysiological correlates of incidentally learned expectations in human vision. Journal of Neurophysiology, 119(4), 1461–1470.

Henson, R.N., Rylands, A., Ross, E., Vuilleumeir, P., & Rugg, M.D. (2004). The effect of repetition lag on electrophysiological and haemodynamic correlates of visual object priming. Neuroimage, 21(4), 1674–1689.

Henson, R. N. (2016). Repetition suppression to faces in the fusiform face area: A personal and dynamic journey. Cortex, 80, 174–184.

Jacobsen, T., & Schroger, E. (2001). Is there pre-attentive memory-based comparison of pitch? Psychophysiology, 38(4), 723–727.

Javitt, D.C., Grochowski, S., Shelley, A-M., & Ritter, W. (1998). Impaired mismatch negativity (MMN) generation in schizophrenia as a function of stimulus deviance, probability, and interstimulus/interdeviant interval. Clinical Neurophysiology, 108, 143–153.

Kaliukhovich, D. A., & Vogels, R. (2016). Divisive normalization predicts adaptation-induced response changes in macaque inferior temporal cortex. Journal of Neuroscience, 36(22), 6116–6128.

Kimura, M., Katayama, J., Ohira, H., & Schroger, E. (2009). Visual mismatch negativity: New evidence from the equiprobable paradigm. Psychophysiology, 46(2), 402–409.

Kimura, M., Schroger, E., & Czigler, I. (2011). Visual mismatch negativity and its importance in visual cognitive sciences. Neuroreport, 22(14), 669–673.

Kleiner, M., Brainard, D., Pelli, D., Ingling, A., Murray, R., & Broussard, C. (2007). What’ s new in Psychtoolbox-3. Perception, 36(14), 1.

Kok, P., Mostert, P., & de Lange, F.P. (2017). Prior expectations induce prestimulus sensory templates. Proceedings of the National Academy of Sciences of the U.S.A., 114(39), 10473–10478.

Kovacs, G., & Vogels, R. (2014). When does repetition suppression depend on repetition probability? Frontiers in Human Neuroscience, 8, 685.

Kremlacek, J., Kreegipuu, K., Tales, A., Astikainen, P., Poldver, N., Naatanen, R., & Stefanics, G. (2016). Visual mismatch negativity (vMMN): A review and meta¬analysis of studies in psychiatric and neurological disorders. Cortex, 80, 76–112.

Kriegeskorte, N., Simmons, W. K., Bellgowan, P. S., & Baker, C. I. (2009). Circular analysis in systems neuroscience: The danger of double dipping. Nature Neuroscience, 12, 535–540.

Laguesse, R., & Rossion, B. (2013). Face perception is whole or none: Disentangling the role of spatial contiguity and interfeature distances in the composite face illusion. Perception, 42(10), 1013–1026.

Leshinskaya, A., & Thompson-Schill, S.L. (2020). Transformation of event representations along middle temporal gyrus. Cerebral Cortex, 30(5), 3148–3166.

Liu-Shuang, J., Norcia, A. M., & Rossion, B. (2014). An objective index of individual face discrimination in the right occipito-temporal cortex by means of fast periodic oddball stimulation. Neuropsychologia, 52, 57–72.

Liu-Shuang, J., Torfs, K., & Rossion, B. (2016). An objective electrophysiological marker of face individualisation impairment in acquired prosopagnosia with fast periodic visual stimulation. Neuropsychologia, 83, 100–113.

May, P. J., & Tiitinen, H. (2010). Mismatch negativity (MMN), the deviance-elicited auditory deflection, explained. Psychophysiology, 47(1), 66–122.

Maheu, M., Dehaene, S., & Meyniel, F. (2019). Brain signatures of a multiscale process of sequence learning in humans. eLife, 2019, 8:e41541.

Maris, E., & Oostenveld, R. (2007). Nonparametric statistical testing of EEG- and MEG-data. Journal of Neuroscience Methods, 164, 177–190.

Movshon, J. A., & Lennie, P. (1979). Pattern-selective adaptation in visual cortical neurones. Nature, 278(5707), 850–852.

Naatanen, R., Gaillard, A. W., & Mantysalo, S. (1978). Early selective-attention effect on evoked potential reinterpreted. Acta Psychologica, 42(4), 313–329.

Nelken, I., & Ulanovsky, N. (2007). Mismatch negativity and stimulus-specific adaptation in animal models. Journal of Psychophysiology, 21(3-4), 214–223.

Pajani, A., Kouider, S., Roux, P., & de Gardelle, V. (2017). Unsuppressable repetition suppression and exemplar-specific expectation suppression in the fusiform face area. Scientific Reports, 7(160).

Patterson, C. A., Wissig, S. C., & Kohn, A. (2013). Distinct effects of brief and prolonged adaptation on orientation tuning in primary visual cortex. Journal of Neuroscience, 33(2), 532–543.

Pincze, Z., Lakatos, P., Rajkai, C., Ulbert, I., & Karmos, G. (2002). Effect of deviant probability and interstimulus/interdeviant interval on the auditory N1 and mismatch negativity in the cat auditory cortex. Cognitive Brain Research, 13, 249–253.

Polich, J., & Margala, C. (1997). P300 and probability: Comparisons of oddball and single-stimulus paradigms. International Journal of Psychophysiology, 25, 169–176.

Quek, G.L., & Rossion, B. (2017). Category-selective human brain processes elicitied in fast periodic visual stimulation streams are immune to temporal predictability. Neuropsychologia, 104, 182–200.

Richter, D., & de Lange, F.P. (2019). Statistical learning attenuates visual activity only for attended stimuli. eLife, 8:e47869

Robinson, J.E., Woods, W., Leung, S., Kaufman, J., Breakspear, M., Young, A.W., & Johnston, P.J. (2019). Prediction-error signals to violated expectations about person identity and head orientation are doubly-dissociated across dorsal and ventral visual stream regions. Neuroimage, 206, 116325.

Rossion, B., & Boremanse, A. (2011). Robust sensitivity to facial identity in the right human occipito-temporal cortex as revealed by steady-state visual-evoked potentials. J Vis, 11(2).

Rostalski, S-M, Amado, C., Kovacs, G., & Feuerriegel, D. (in press). Measures of repetition suppression in the Fusiform Face Area are inflated by co-occurring effects of statistically learned visual associations. BiorXiv Preprint. doi https://doi.org/10.1101/803163

Sawamura, H., Orban, G.A., & Vogels, R. (2006). Selectivity of neuronal adaptation does not match response selectivity: A single-cell study of the fMRI adaptation paradigm. Neuron, 49, 307–318.

Schomaker, J., & Meeter, M. (2015). Short- and long-lasting consequences of novelty, deviance and surprise on brain and cognition. Neuroscience and Biobehavioural Reviews, 55, 268–279.

Schultz, W. (2016). Dopamine reward prediction-error signalling: A two-component response. Nature Reviews Neuroscience, 17, 183–195.

Smout, C., Garrido, M.I., & Mattingley, J.B. (2020). Global effects of feature-based attention depend on surprise. Neuroimage, 215, 116785.

Solomon, S. G., & Kohn, A. (2014). Moving sensory adaptation beyond suppressive effects in single neurons. Current Biology, 24(20), R1012–1022.

Stefanics, G., Kimura, M., & Czigler, I. (2011). Visual mismatch negativity reveals automatic detection of sequential regularity violation. Frontiers in Human Neuroscience, 5, 46.

Stefanics, G., Astikainen, P., & Czigler, I. (2014). Visual mismatch negativity (vMMN): A prediction error signal in the visual modality. Frontiers in Human Neuroscience, 8, 1074.

Stefanics, G., Heinzle, J., Horvath, A.A., & Stephan, K.E. (2018). Visual mismatch negativity and predictive coding: A computational single-trial ERP study. The Journal of Neuroscience, 38(16), 4020–4030.

Stefanics, G., Stephan, K.E., & Heinzle, J. (2019). Feature-specific prediction errors for visual mismatch. Neuroimage, 196, 142–151.

Summerfield, C., Trittschuh, E. H., Monti, J. M., Mesulam, M. M., & Egner, T. (2008). Neural repetition suppression reflects fulfilled perceptual expectations. Nature Neuroscience, 11(9), 1004–1006.

Summerfield, C., Wyart, V., Johnen, V. M., & de Gardelle, V. (2011). Human scalp electroencephalography reveals that repetition suppression varies with expectation. Frontiers in Human Neuroscience, 5, 1–13.

Todorovic, A., & de Lange, F. P. (2012). Repetition suppression and expectation suppression are dissociable in time in early auditory evoked fields. Journal of Neuroscience, 32(39), 13389–13395.

Ulanovsky, N., Las, L., Farkas, D., & Nelken, I. (2004). Multiple time scales of adaptation in auditory cortex neurons. The Journal of Neuroscience, 24(46), 10440–10453.

Vinken, K., & Vogels, R. (2017). Adaptation can explain evidence for encoding of probablistic information in macaque inferior temporal cortex. Current Biology, 27, R1210–R1211.

Wacongne, C., Labyt, E., van Wassenhove, V., Bekinschtein, T., Naccache, L., & Dehaene, S. (2011). Evidence for a hierarchy of predictions and prediction errors in human cortex. Proceedings of the National Academy of Sciences of the United States of America, 108(51), 20754–20759.

Wessel, J. R. (2017). An adaptive orienting theory of error processing. Psychophysiology, 55, 1–21.

Wessel, J.R., & Huber, D. (2019). Frontal cortex tracks surprise separately for different sensory modalities but engages in a common inhibitory control mechanism. PLoS Computational Biology, 15(7), e1006927. doi:10.1371/journal.pcbi.1006927

Willenbockel, V., Sadr, J., Fiset, D., Horne, G. O., Gosselin, F., & Tanaka, J. W. (2010). Controlling low-level image properties: The SHINE toolbox. Behavior Research Methods, 42(3), 671–684.

