## Supplementary Material for "Visual mismatch responses index surprise signalling but not expectation suppression"

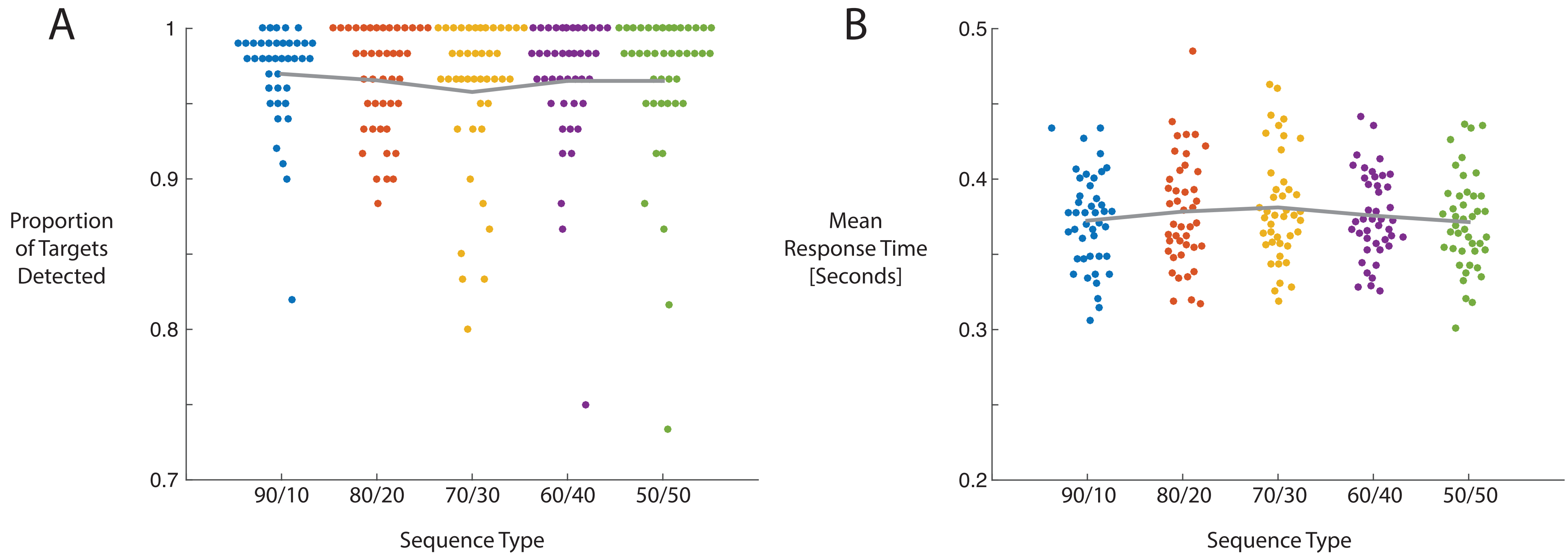


**Supplementary Figure S1**. Performance on the fixation cross colour change detection task for each participant, split by sequence type. A) Proportions of correctly identified targets. B) Mean RTs for correctly identified targets. Grey lines depict group means for each sequence type.

**Supplementary Table S1**. *Summary statistics for numbers of epochs retained for analyses per participant, split by stimulus appearance probability condition.*

| Oddball Appearance Probability | Mean | Median | SD | Min | Max |
|---|---|---|---|---|---|
| 10% | 67 | 68 | 4 | 54 | 76 |
| 20% | 81 | 82 | 4 | 70 | 97 |
| 30% | 121 | 124 | 7 | 102 | 144 |
| 40% | 162 | 165 | 8 | 119 | 168 |
| 50% | 402 | 409 | 20 | 318 | 419 |
| 60% | 243 | 246 | 12 | 180 | 252 |
| 70% | 283 | 288 | 18 | 216 | 338 |
| 80% | 324 | 330 | 19 | 272 | 388 |
| 90% | 606 | 617 | 34 | 508 | 688 |

**Supplementary Table S2**. *Summary statistics for numbers of epochs retained for analyses per participant, after excluding epochs corresponding to oddballs that immediately follow a surprising oddball.*

| Oddball Appearance Probability | Mean | Median | SD | Min | Max |
|---|---|---|---|---|---|
| 10% | 67 | 68 | 4 | 54 | 76 |
| 20% | 71 | 73 | 4 | 63 | 87 |
| 30% | 93 | 94 | 6 | 76 | 108 |
| 40% | 109 | 112 | 7 | 79 | 117 |
| 50% | 402 | 409 | 20 | 318 | 419 |
| 60% | 132 | 133 | 8 | 105 | 142 |
| 70% | 186 | 189 | 13 | 139 | 224 |
| 80% | 249 | 253 | 15 | 209 | 298 |
| 90% | 531 | 540 | 29 | 445 | 602 |

**Supplementary Table S3**. *Results of mean amplitude comparisons between each oddball probability condition and the 50% probability (expectation neutral) oddball condition, for the early (200-350 ms) time window. Statistically significant (p < 0.05) results are denoted by bold text. CI = Frequentist 95% Confidence Interval.*

| Oddball Appearance Probability | df | t | p | $BF_{10}$ | Mean Difference [μV] | CI Lower Bound | CI Upper Bound |
|---|---|---|---|---|---|---|---|
| Surprising (10-40%) | 42 | -2.94 | **0.005** | 6.78 | -0.33 | -0.56 | -0.10 |
| 10% | 42 | -2.25 | **0.030** | 1.59 | -0.39 | -0.74 | -0.04 |
| 20% | 42 | -1.73 | 0.092 | 0.64 | -0.31 | -0.68 | 0.05 |
| 30% | 42 | -2.04 | **0.048** | 1.07 | -0.26 | -0.52 | -0.002 |
| 40% | 42 | -2.94 | **0.005** | 6.83 | -0.35 | -0.59 | -0.11 |
| Expected (60-90%) | 42 | -0.79 | 0.433 | 0.22 | -0.07 | -0.25 | 0.10 |
| 60% | 42 | -2.19 | **0.035** | 1.40 | -0.21 | -0.40 | -0.02 |
| 70% | 42 | -0.19 | 0.853 | 0.17 | -0.02 | -0.26 | 0.21 |
| 80% | 42 | -1.14 | 0.259 | 0.30 | -0.13 | -0.36 | 0.10 |
| 90% | 42 | 0.77 | 0.448 | 0.22 | 0.08 | -0.13 | 0.28 |

**Supplementary Table S4**. *Results of mean amplitude comparisons between each oddball probability condition and the 50% probability (expectation neutral) oddball condition, for the late (500-1,000 ms) time window. Statistically significant ($p < 0.05$) results are denoted by bold text. CI = Frequentist 95% Confidence Interval.*

| Oddball Appearance Probability | df | t | p | $BF_{10}$ | Mean Difference [μV] | CI Lower Bound | CI Upper Bound |
|---|---|---|---|---|---|---|---|
| Surprising (10-40%) | 42 | -4.84 | **< 0.001** | 1141.24 | -0.27 | -0.39 | -0.16 |
| 10% | 42 | -3.32 | **0.002** | 17.27 | -0.40 | -0.64 | -0.16 |
| 20% | 42 | -1.92 | 0.062 | 0.87 | -0.29 | -0.47 | 0.01 |
| 30% | 42 | -2.87 | **0.006** | 5.87 | -0.27 | -0.45 | -0.08 |
| 40% | 42 | -2.59 | **0.013** | 3.17 | -0.20 | -0.36 | -0.05 |
| Expected (60-90%) | 42 | -0.04 | 0.966 | 0.17 | -0.002 | -0.11 | 0.10 |
| 60% | 42 | -0.06 | 0.952 | 0.17 | -0.01 | -0.19 | 0.18 |
| 70% | 42 | 0.08 | 0.934 | 0.17 | 0.01 | -0.14 | 0.15 |
| 80% | 42 | -0.48 | 0.633 | 0.18 | -0.03 | -0.15 | 0.09 |
| 90% | 42 | 0.42 | 0.674 | 0.18 | 0.02 | -0.08 | 0.12 |

### Mass Univariate Analyses of Contrasts Between Surprising, Neutral and Expected Conditions

Here we have plotted the results of mass-univariate analyses using paired-samples t tests to compare surprising and expected conditions, which were used to derive the early and late time window ROIs for further analyses. Methods for this analysis are detailed in section 2.8 of the manuscript. Briefly, we ran paired-samples t tests to compare surprising and expected conditions (alpha level 0.05), and defined our ROIs based on the time windows during which surprising oddballs evoked more negative-going amplitudes at posterior electrodes. We have also plotted the results after controlling the false discovery rate (FDR) at $q = 0.05$ using the Benjamini-Hochberg procedure. In addition, we have plotted the results of [surprising – neutral] and [expected – neutral] contrasts, however the results of these analyses were not considered during definition of the ROIs. These plots show that the spatiotemporal pattern of [surprising – neutral] differences is similar to the pattern of [surprising – expected] amplitude differences. By contrast, the sparse pattern of statistically significant [expected – neutral] differences does not resemble patterns of real effects as found when measuring ERPs, and did not survive FDR correction.

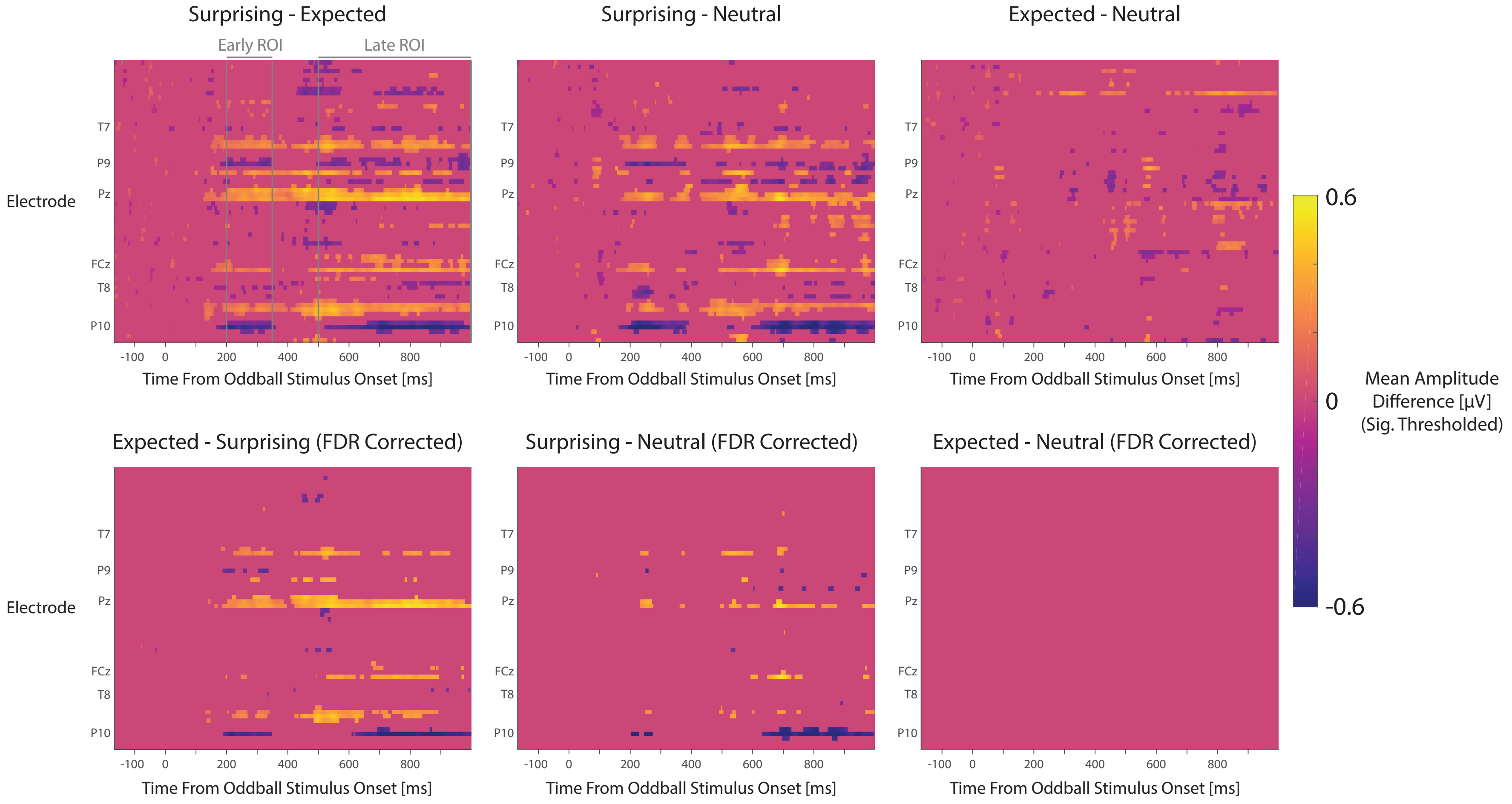


**Supplementary Figure S2.** Channel-by-timepoint matrices of group mean amplitude differences (thresholded by statistical significance, alpha = 0.05) for [surprising – expected], [surprising – neutral] and [expected – neutral] contrasts. Results are shown for analyses uncorrected for multiple comparisons (top row) and when controlling the false discovery rate (FDR) at q = 0.05 using the Benjamini-Hochberg procedure (bottom row). The time windows of the early (200-350 ms) and late (500-1000 ms) ROIs are marked in the plot of surprising – expected differences.

### Results of Analyses Using Positive-Going [Surprise – Expected] Difference ROIs

The following sections describe the results of analyses using ROIs defined by more positive-going ERPs following surprising compared to expected oddball stimuli, at electrodes POz, PO3, Pz, P1, P2, P4, CP3, CPz, CP1, CP2, Cz, FCz, and FC2. Group-averaged ERPs at these electrodes for expected, neutral and surprising conditions are plotted in Supplementary Figure S2.

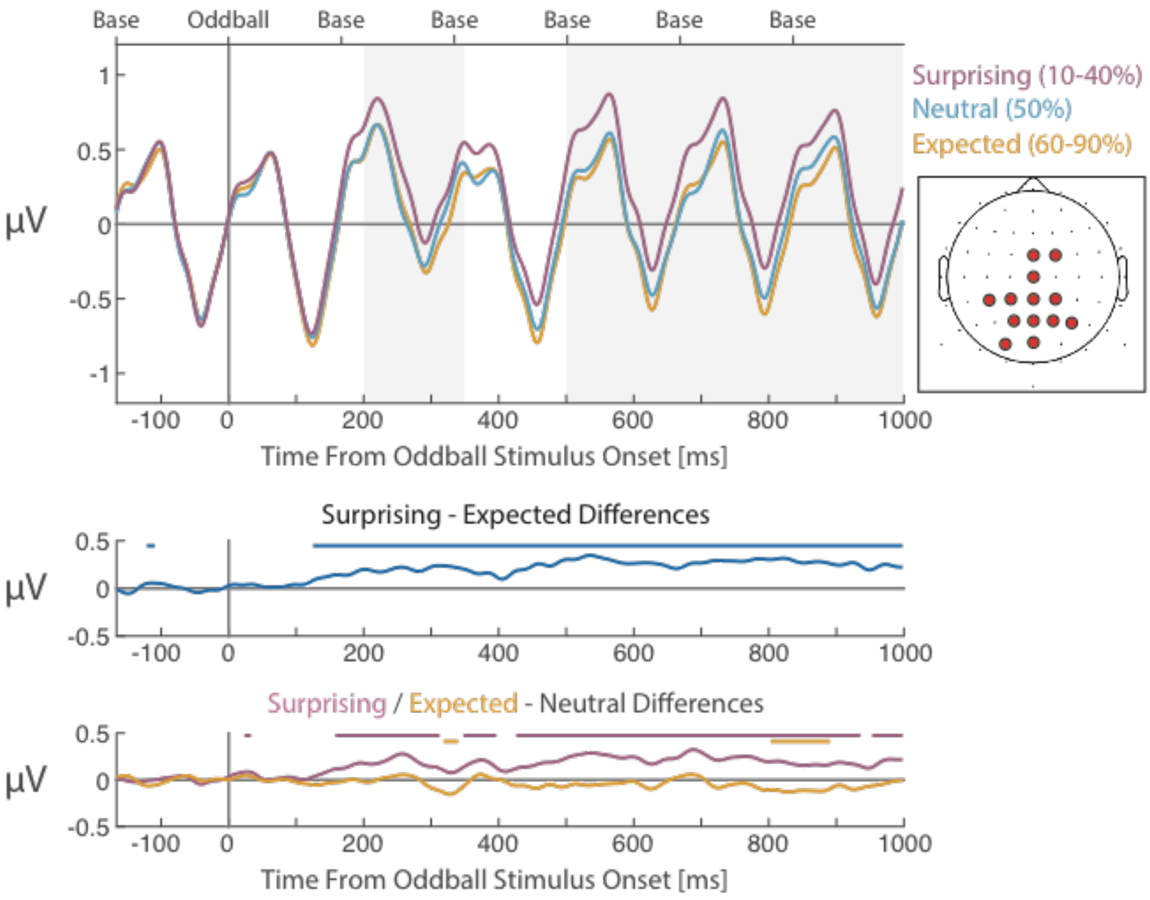


**Supplementary Figure S3**. Group-averaged ERPs evoked by surprising (10-40% probability), neutral (50%) and expected (60-90%) oddballs. Group-averaged ERPs for each condition (top panel), differences between surprising and expected conditions (middle panel), and differences between surprising and expected conditions compared to the neutral condition (bottom panel), averaged across electrodes POz/3, Pz/1/2/4, CPz/1/2/3, Cz, and FCz/2 (electrode locations shown in the inset scalp map). Grey shaded areas denote the early (200-350 ms) and late (500-1000 ms) ROI time windows. Labels at the top of the upper panel denote the onset times of the base and oddball stimuli within the epoch. Thick lines at the top of each plot denote statistically significant differences between ERPs that were averaged over selected electrodes ($p < .05$ uncorrected). Note that this figure depicts ERPs calculated after removal of epochs immediately following a surprising oddball.

**Quantifying Effects of Fulfilled Expectations and Surprise**

To isolate and quantify effects of expectation suppression and surprise, we compared mean amplitudes for expected and surprising oddballs with those for neutral oddballs. Results of paired-samples t tests are displayed in Supplementary Table S5 for the early time window, and Supplementary Table S6 for the late time window. Mean amplitudes were more positive-going for surprising compared to neutral oddballs, for the early time window ($p < 0.001$, $BF_{10} = 65.83$) and the late time window ($p < 0.001$, $BF_{10} = 1722.77$). By contrast, amplitudes did not appear to differ between expected and neutral conditions, with Bayes factors indicating evidence for the null hypothesis for the early time window, and only a small amount of evidence favouring the null for the late time window (early time window: $p = 0.487$, $BF_{10} = 0.21$, late time window: $p = 0.137$, $BF_{10} = 0.48$).

We next compared the neutral condition with each stimulus appearance probability condition to assess whether this pattern of results was consistent with the condition-averaged data. Scatterplots of mean amplitudes by stimulus probability are displayed in Supplementary Figures S3A (early time window) and S3B (late time window). Relevant statistical outputs are summarised in Supplementary Tables S5 and S6. ROI-averaged amplitudes were more negative-going for all surprising oddball conditions across both time windows (p's < 0.05) except for the 20% appearance probability condition (early time window $p = 0.297$, late time window $p = 0.054$). Group mean [surprising – neutral] difference magnitudes were similar across all surprising oddball conditions, ranging from -0.10 to 0.25 μV (early time window) and 0.18 to 0.29 μV (late time window). By contrast, none of the expected oddball conditions evoked significantly more positive or negative amplitudes compared to the neutral condition (all p's > 0.077). Group mean [expected – neutral] difference magnitudes were numerically smaller than the observed surprise effects, and ranged from -0.09 to 0.02 μV (early time window) and -0.11 to 0.01 μV (late time window).

**Graded Effects of Stimulus Appearance Probability**

After identifying contributions of surprise to VMR amplitudes, we then assessed whether there were graded differences in ROI-averaged amplitudes by oddball presentation probability for surprising (10-40%) oddballs. For the early time window beta values did not significantly differ from zero, and the Bayes factor indicated moderate evidence for the null hypothesis, $t(42) = -0.13$, $p = 0.895$, $BF_{10} = 0.17$, mean beta value -0.0004, CI [-0.01, 0.01]. Here, the mean estimated beta value denotes an average change of only -0.0004 μV for each percentage increase in oddball appearance probability. We found similar results for the late time window, $t(42) = 0.09$, $p = 0.927$, $BF_{10} = 0.17$, mean beta value 0.0003, CI [-0.01, 0.01].

**Supplementary Table S5**. *Results of mean amplitude comparisons between each oddball probability condition and the 50% probability (expectation neutral) oddball condition, for the early (200-350 ms) time window. Statistically significant (p < 0.05) results are denoted by bold text. CI = Frequentist 95% Confidence Interval.*

| Oddball Appearance Probability | df | t | p | $BF_{10}$ | Mean Difference [μV] | CI Lower Bound | CI Upper Bound |
|---|---|---|---|---|---|---|---|
| Surprising (10-40%) | 42 | 3.84 | **< 0.001** | 65.83 | 0.17 | 0.08 | 0.27 |
| 10% | 42 | 3.26 | **0.002** | 14.74 | 0.25 | 0.09 | 0.40 |
| 20% | 42 | 1.06 | 0.297 | 0.28 | 0.10 | -0.09 | 0.29 |
| 30% | 42 | 2.48 | **0.017** | 2.50 | 0.15 | 0.03 | 0.28 |
| 40% | 42 | 3.00 | **0.005** | 7.91 | 0.20 | 0.06 | 0.33 |
| Expected (60-90%) | 42 | -0.70 | 0.487 | 0.21 | -0.03 | -0.13 | 0.06 |
| 60% | 42 | 0.22 | 0.828 | 0.17 | 0.02 | -0.14 | 0.17 |
| 70% | 42 | -1.49 | 0.143 | 0.46 | -0.09 | -0.22 | 0.03 |
| 80% | 42 | 0.30 | 0.767 | 0.17 | 0.02 | -0.11 | 0.14 |
| 90% | 42 | -1.30 | 0.201 | 0.36 | -0.08 | -0.20 | 0.04 |

**Supplementary Table S6**. *Results of mean amplitude comparisons between each oddball probability condition and the 50% probability (expectation neutral) oddball condition, for the late (500-1000 ms) time window. Statistically significant (p < 0.05) results are denoted by bold text. CI = Frequentist 95% Confidence Interval.*

| Oddball Appearance Probability | df | t | p | $BF_{10}$ | Mean Difference [μV] | CI Lower Bound | CI Upper Bound |
|---|---|---|---|---|---|---|---|
| Surprising (10-40%) | 42 | 4.97 | < **0.001** | 1722.77 | 0.22 | 0.13 | 0.30 |
| 10% | 42 | 2.24 | **0.031** | 1.55 | 0.18 | 0.02 | 0.34 |
| 20% | 42 | 1.98 | 0.054 | 0.98 | 0.21 | -0.004 | 0.43 |
| 30% | 42 | 4.27 | < **0.001** | 217.76 | 0.29 | 0.16 | 0.43 |
| 40% | 42 | 2.46 | **0.018** | 2.42 | 0.18 | 0.03 | 0.33 |
| Expected (60-90%) | 42 | -1.52 | 0.137 | 0.48 | -0.06 | -0.13 | 0.02 |
| 60% | 42 | -1.19 | 0.239 | 0.32 | -0.07 | -0.20 | 0.05 |
| 70% | 42 | -1.81 | 0.078 | 0.73 | -0.11 | -0.23 | 0.01 |
| 80% | 42 | -1.31 | 0.198 | 0.37 | -0.06 | -0.15 | 0.03 |
| 90% | 42 | 0.22 | 0.828 | 0.17 | 0.01 | -0.07 | 0.09 |

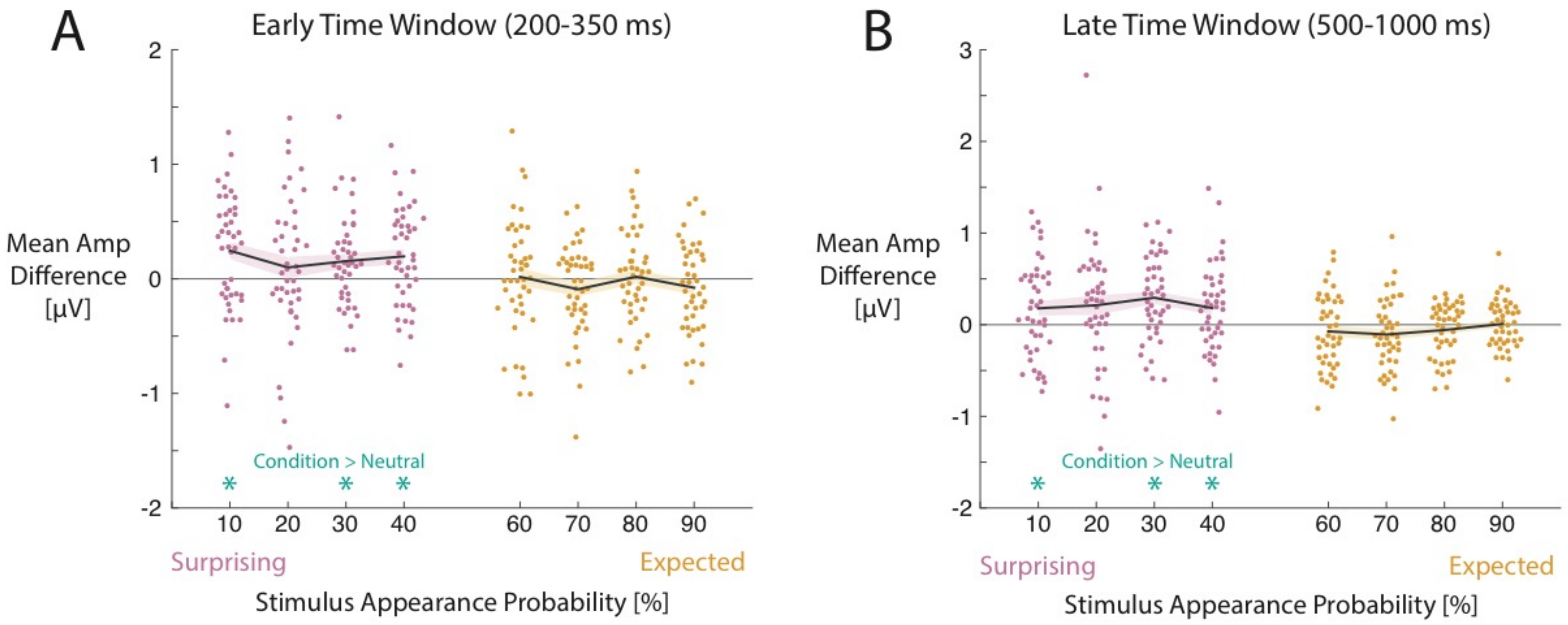


**Supplementary Figure S4**. Mean amplitudes for A) early (200-350 ms) and B) late (500-1000 ms) time windows, sorted by stimulus appearance probability. Thick black lines represent the group mean differences. Shaded regions represent standard errors. Asterisks at the bottom of each plot indicate statistically significant ($p < 0.05$) differences.
